# Trophic and temporal dynamics of macrophage biology in human inner ear organogenesis

**DOI:** 10.1101/2025.05.16.654631

**Authors:** Yidi Deng, Boaz Ehiogu, Emilia Luca, Alain Dabdoub, Kim-Anh Lê Cao, Christine A. Wells, Bryony A. Nayagam

## Abstract

Recent single-cell transcriptomic approaches are uncovering the breadth and depth of cell diversity within the mammalian inner ear. Macrophages, detected from gestational week 7 in the human inner ear, persist into adulthood, and yet remain poorly understood in terms of their origin and function. Using self-generated and public scRNA-seq data, we identify seven distinct macrophage subtypes spanning fetal weeks 7.5 to 16.4 and adulthood. Each macrophage subtype is linked to specific developmental stages and displays a unique gene expression profile. These findings corroborate earlier histological evidence of resident and non-resident macrophages in both the developing and adult human cochlea. We also show that the human inner ear is seeded by macrophages from both embryonic and more definitive sources, corroborating studies in mouse. By analyzing ligand-receptor interactions, we highlight potential macrophage contributions to inner ear organogenesis. This research provides new insights into the diverse roles of human inner ear macrophages.

## 1 Introduction

Normal development of the inner ear requires a sophisticated orchestration of specialized cell differentiation and integration, ultimately giving rise to the exquisite organs of hearing and balance. However, the timeline and dynamic nature of this developmental process remain only partially understood. Recent single-cell transcriptomic studies have improved our understanding of the molecular phenotypes of the mammalian inner ear through differential gene expression analyses in both mouse [Kolla et al., 2020, Petitpŕe et al., 2022] and human [van der Valk et al., 2023]. While these studies have focused primarily on the inner ear hair cells and neurons, there are at least 17 other cell types present in developing human inner ear (HIE), including large numbers of mesenchymal cells, supporting cells, and also macrophages [van der Valk et al., 2023]. Each cell types likely performs a specific, but as yet largely uncharacterized, function in the formation of this elaborate organ.

Inner ear macrophages (IEMs) have gained increasing attention for their innate and adaptive immune roles in the normal [Liu and Rask-Andersen, 2019], noise-damaged [He et al., 2020], and cochlear-implanted [Bas et al., 2015, Nadol Jr et al., 2014] HIE. In mouse, they have also been implicated in repairing the utricle [Kaur et al., 2015a] and in protecting cochlear afferent neurons [Kaur et al., 2015b]. During HIE development, macrophages have been first detected at gestational week 7, marked by the expression of *IBA1* and *CD45* [Steinacher et al., 2022], and they clearly populate in the adult cochlea [Liu and Rask-Andersen, 2019]. Yet, little is known about the origin of human cochlear macrophages or their functional contributions to inner ear organogenesis.

Beyond traditional immune surveillance, macrophages have well-established roles in development and tissue homeostasis [Mosser et al., 2021, Wynn et al., 2013]. They infiltrate numerous organs in the developing human [Bian et al., 2020], where they perform diverse and critical functions for normal organogenesis. For instance, in the brain, macrophages have been shown to regulate neurogenesis, synaptic pruning, and the clearance of apoptotic cells during brain development, thereby shaping the intricate neural circuits fundamental to normal function [Paolicelli et al., 2011]. In the retina, macrophages help pattern and vascularize developing tissue, ultimately ensuring proper pupil morphology [Takahashi et al., 2020]. Similarly, in the lung, macrophages assist in alveolar development and surfactant homeostasis, facilitating proper respiratory function [Schneider et al., 2014]. Given the complexity of inner ear patterning, fluid homeostasis and vascularization, along with the observed early presence of macrophages during inner ear development, these multifunctional cells may be key to better understanding the intricate processes of normal auditory and vestibular organ formation.

In the present study, we offer the first comprehensive overview of macrophage molecular heterogeneity in the HIE (Figure 1A). By integrating both public and newly generated transcriptomic datasets, we assembled a comprehensive macrophage atlas covering key stages of human inner ear (HIE) development—from early fetal weeks (FWs) 7.5 and 9.2, to middle FWs 16 and 16.4, and through to adulthood. We identify several distinct transcriptional profiles that define trophic roles for IEMs at different developmental ages and predict their likely modes of communication with other cell types. We also compared IEM phenotypes with macrophages present in other tissues during a similar window of human development [Bian et al., 2020]. Collectively, these findings provide essential insights into the diverse roles of IEMs during HIE development and pave the way for macrophage-targeted strategies to prevent or treat inner ear disorders.

**Figure 1.**
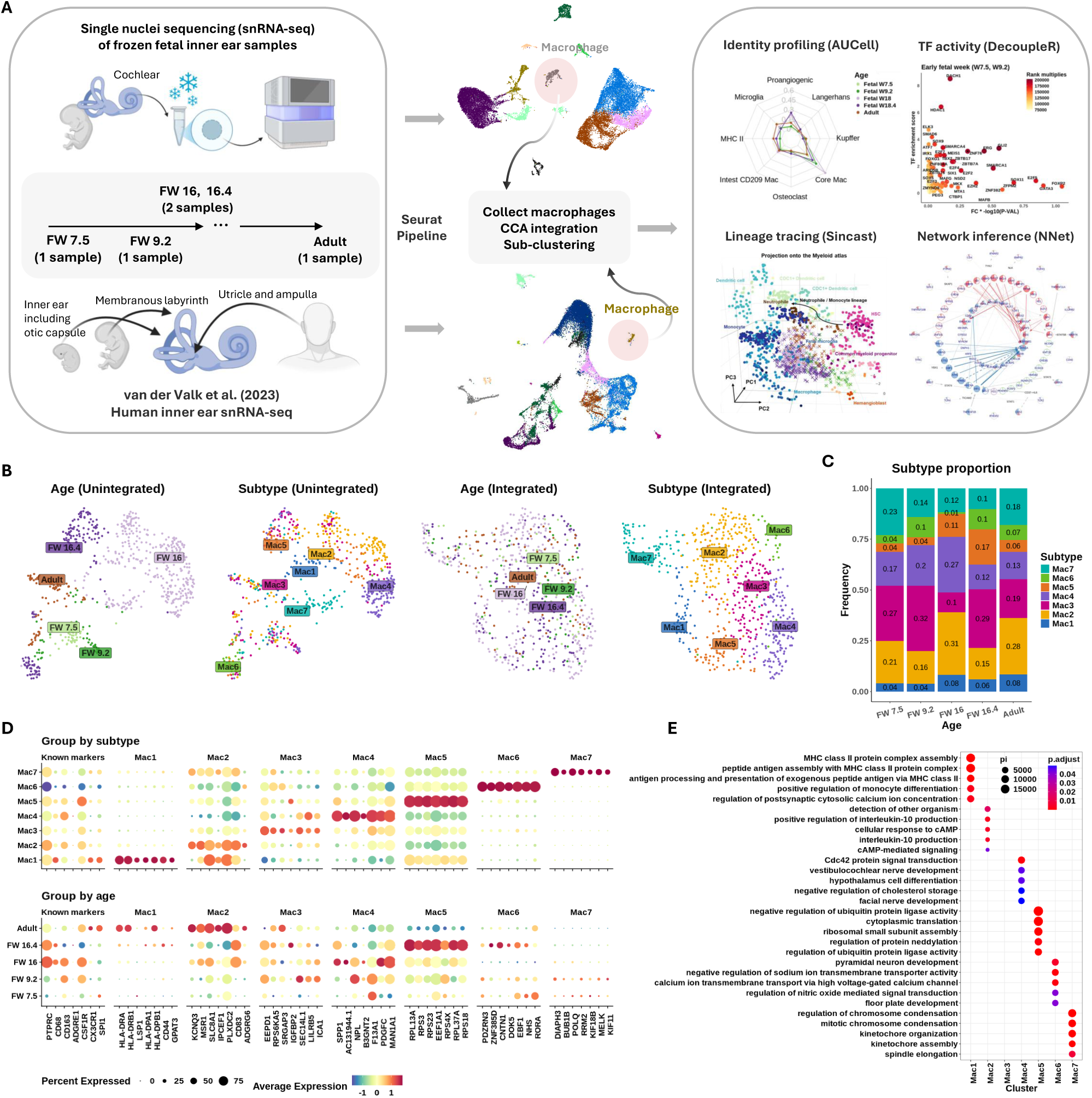
Seven fetal macrophage subtypes found in the developing inner ear. **(A)** Schematic of (L-R) tissue sample, integration and analysis workflows used in this study. **(B)** UMAP plots showing grouping of cells before (L) and after (R) Seurat integration. Macrophage subtypes were identified using Leiden clustering. **(C)** Stacked bar plot showing proportions (y-axis) of each subtype across developmental age (x-axis). **(D)** Expression of common macrophage markers, and top ranked discriminating genes (first 7 genes) in each subtype. y-axis shows categories of subtype (top) and age (bottom). Gene symbols shown on x-axis. Column headers indicate the subtypes that the markers represent. Expression shown on column-normalized z-score (red highest expression, blue lowest, yellow indicating mean expression value). The size of the dot indicates the percentage of the group expressing the gene of interest. **(E)** Gene set over-representation analysis of genes differentially expressed between macrophage subtypes. Subtypes shown on x-axis, Gene ontology pathway terms on y axis. Enrichment shown as circle size (relative proportion) and colour (adjusted p-value).

## 2 Results

### 2.1 There are seven distinct macrophage subtypes present in the human inner ear

The HIE was found to contain seven distinct macrophage subtypes (denoted as Mac 1 to Mac 7) each with a unique transcriptomic profile (Figure 1B,D). All subtypes were present throughout inner ear development (Figure 1C), noting that middle FW samples (FWs 16 and 16.4) were dissected slightly differently to those at early FWs (FWs 7.5 and 9.2) and adult tissue. Middle FW tissues yielded macrophages only from the cochlear modiolus rather than from the entire inner ear. Further analysis showed that Mac 3 likely represents an intermediate cell state between Mac 2 and 4, given the small number of differentially expressed genes (DEGs) that distinguish it (Supplementary Figure S1C,D).

Previous studies using immunohistochemistry to identify macrophages in the HIE have reported the expression of *PTPRC* (CD45), *CD68*, *CD163* and *CX3CL1* [O’Malley et al., 2016, Liu and Rask-Andersen, 2019, Steinacher et al., 2022]. Consistently, we confirmed the expression of these markers (along with additional selected known macrophage markers *CSFR1* and *SPI1*) in both fetal and adult IEMs. However, these markers were non-discriminatory in defining any particular macrophage subtype (Figure 1D “known marker”). Note that we isolated the macrophage subset of the inner ear data based on the expression of *PTPRC* and *ITGAM* (CD11b), which are commonly used markers for gating macrophages (Supplementary Figure S1A,B). In contrast, *ADGRE1* (F4/80) expression was absent in our samples, in agreement with known species-specific expression patterns [Waddell et al., 2018], and its enrichment in human eosinophils [Hamann et al., 2007] (Figure 1D).

Differential expression (DE) analysis revealed seven distinct phenotypes, suggesting unique functional roles in early development (Mac 6 and 7), trophic support, and immune homeostasis (Mac 3, 4, and 5), and antigen presentation (Mac 1 and 2). The DE markers of Mac 5, 6, and 7 were distinctly expressed in their defining subtypes across all developmental stages, whereas those of Mac 1, 2, and 3 formed distinct patterns mostly after FW 16 (Supplementary Figure S2).

Specifically, Mac 7 populations were enriched for genes implicated in important roles in cell proliferation (*MELK*), cell division (*BUB1* and *KIF11*), DNA synthesis and repair (*POLQ* and *RRM2*), and cytoskeletal remodeling (*DIAPH3*) [Whitfield et al., 2006, Locard-Paulet et al., 2022]. Mac 6 was observed to have additional regulatory functions, expressing contactin 1 (*CNTN1*) [Chatterjee et al., 2019], regulators of Wnt signaling (*DOK5* and *PDZRN3*) [Xu et al., 2022, Wu et al., 2021], and the nuclear hormone receptor (*RORA*) [Cheng et al., 2024], all of which have been implicated in cell proliferation events (Figure 1E). Notably, Wnt signaling plays a critical role in inner ear development and function [Dabdoub et al., 2003, Geng et al., 2016, Munnamalai and Fekete, 2013]. Further analysis of selected growth factor expression revealed that this macrophage subtype was involved in both neuregulin and Wnt5 signaling during early developmental ages (Supplementary Figure S3).

By contrast, Mac 3, 4, and 5 displayed broader associations with facial and vestibulocochlear nerve development (Figure 1E). For instance, Mac 3 and 5 indicated a trophic phenotype through their relative expression of *IGFBP2* (Mac 3) and *PDGF* (Mac 5), respectively. These growth factors are vital for both vascular and cochlear neuro-sensory development and preservation. Additionally, Mac 4 showed functional specialization in tissue remodeling, evidenced by the expression of secreted phosphoprotein 1 (*SPP1*), transglutimase *F13A1*, and acetylglucosaminyltransferase *B3GNT2*, which are genes involved in crosslinking and modifying matrix proteins.

Mac 2 and 3 also expressed mediators of efferocytosis, including *MSR1* (a scavenger receptor), *SEC14L1* (a lipid co-factor that inhibits RIG-I signaling [Li et al., 2013]), *EEPD1* (involved in cholesterol efflux), *ICA1* (lipid complexing and receptor trafficking), and *SRGAP3* (a regulator of actin dynamics via *RAC1*). They also expressed regulators of calcium and potassium efflux (*KCNQ3* and *SLC8A1*), along with growth factor-binding complexes such as *IGFBP2*, *PLXCD2* (PEDF binding), and the adhesion G-protein–coupled receptor *ADGRG6*. While Mac 2 markers were predominantly expressed in adult IEMs, Mac 3 markers were enriched in early fetal IEMs at FWs 7.5 and 9.2.

Mac 1 and 2 populations represented a mature, “classical” macrophage phenotype characterized by immune surveillance functions. Their putative roles were supported by the expression of numerous HLA transcripts (Figure 1D), indicating active antigen presentation and MHC-II regulation (Figure 1E). Over-representation analysis of subtype markers further revealed enrichment in sodium and calcium ion transport pathways (Figure 1E), suggesting that Mac 1 and 2 may support hair cell and neural function. In addition, these macrophages may contribute to the maintenance of inner ear fluid (endolymph) homeostasis, given that these cations are critical for the generation of neural action potentials that underpin hearing and balance [Salt et al., 1987]. Mac 1 also exhibited monocyte-like features and was most abundant from FW 16 through adulthood, comprising more than 6% of the population. The relative expression dynamics of each macrophage subtype across developmental time points (Figure 1D) are further examined in Figure 2.

**Figure 2.**
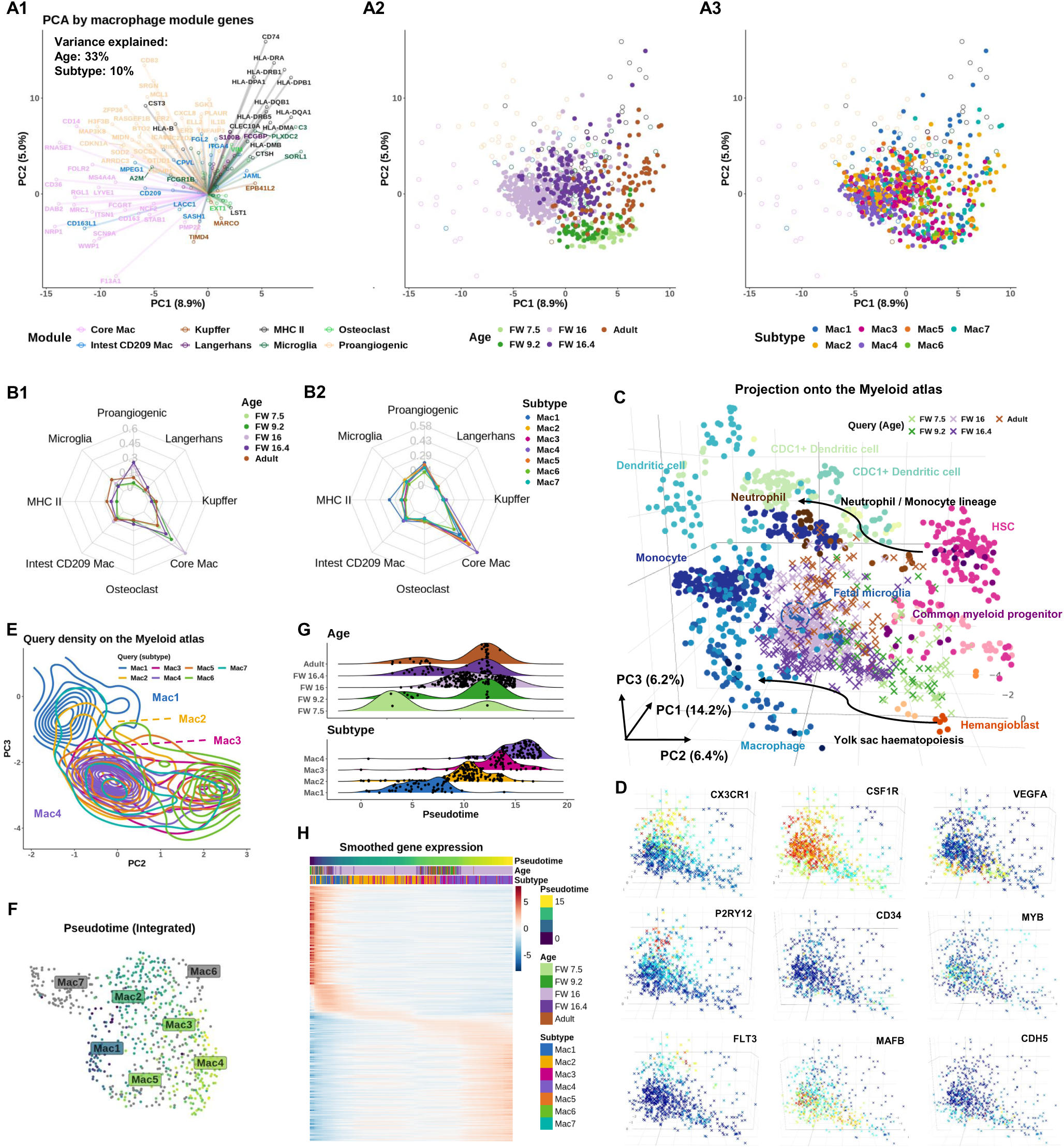
Age-dependent recruitment of macrophages to the inner ear shows defined subtypes present at distinct times. **(A)** Biplot showing PCA based on macrophage module genes [Wang et al., 2023]. (A1) Circles represents gene loadings, colored by tissue module. PC scores of inner ear macrophages (IEMs) denoted as dots, colored by (A2) donor age and (A3) subtypes. **(B)** Radar graphs showing AUCell score of macrophage modules genes at different (B1) developmental stages and (B2) different IEM subtypes in the inner ear **(C)** Projection of IEM subtypes to the Stemformatics.org myeloid atlas. Atlas samples denoted as circles, projected inner ear data as crosses. Inner ear samples coloured by ages. **(D)** Projected IEMs colored by the expression of *CX3CR1* (adult); *CSF1R* (all); VEGFA (fetal weeks, FWs 7.5, 9.2, 16.4); *P2RY12* (FWs 16 and adult); CD34 (FWs 7.5 and 9.2); *MYB* and *FLT3* (FWs 16 and adult); *MAFB* and *CDH5* (FWs 7.5, 9.2, and 16.4). **(E)** Contour map of projected IEMs coloured by subtypes demonstrates a continuum between Mac 1, 2, 3, and 4. **(F)** UMAP plot, colored by pseudotime inferred using b mn,shot with Mac 1 selected as the root, highlights a trajectory of projected IEMs. Mac 5, 6, and 7 are excluded from the trajectory. **(G)** Pseudotime alignment of macrophages shows subtype (bottom plot), rather than age (top plot), is the strongest predictor of pseudotime on the stemformatics atlas. **(H)** A heatmap of the top 500 genes correlated with pseudotime, clustered along the^9^pseudotime axis showing transition from Mac 1 to Mac 4.

### 2.2 Age-dependent recruitment of macrophages to the inner ear shows defined subtypes present at distinct times

Having identified seven macrophage subtypes, we next investigated whether specific macrophage phenotypes were present or absent at different stages of inner ear development. Figure 1C and 1D indicates a possible enrichment of Mac 1 and 2 in adult tissue; Mac 3, 4, and 5 at middle FWs; and Mac 6 and 7 at early FWs. To test this, we performed principal component analysis (PCA) on the expression of established macrophage module genes from Wang et al. [2023] to examine their association with developmental age and IEM subtypes.

The biplot in Figure 2A overlays (i) a loading plot (A1), which highlights the genes that drive variance, with (ii) the projection of individual IEMs in the PCA space (A2–A3). The cells separate clearly by developmental ages, whereas subtype has little influence, suggesting that IEMs acquire distinct module identities during development. FW 16 IEMs displayed a strong core macrophage identity, while adult IEMs exhibited MHC-II and microglial signatures from the opposite side of the PCA. FW 16.4 macrophages occupied an intermediate, pro-angiogenic niche, and early FWs IEMs clustered at the center of the PCA, indicating a relatively immature state. In additional studies, we showed that the population of IEMs that we have analyzed shares the greatest similarity with macrophage populations in the skin and brain during human development (Supplementary Figure S4).

We validated these findings with AUCell [Aibar et al., 2017], which calculates module activity scores per cell in a manner that is robust to batch effects. This confirmed the age-dependent module activities (Figure 2B1). When examining module scores by subtypes, all subtypes exhibited a strong core macrophage identity. Notably, Mac 1 showed the highest co-adoption of MHC-II, microglial, and pro-angiogenic markers, consistent with a mature, efferocytotic, and antigen-presenting phenotype enriched in adult IEMs (Figure 1E).

To further interrogate the effect of age on macrophage identity, we used Sincast [Deng et al., 2022] to project our IEM data (colored by donor ages) onto a myeloid atlas [Rajab et al., 2021] consisting of bulk transcriptomic data from 44 independent studies. This analysis revealed two macrophage developmental trajectories: one from common myeloid progenitors (CMPs) / hematopoetic stem cells (HSCs) and the second from a hemangioblast-like progenitors (HLPs); both of which ultimately differentiated into a fetal microglial-like or mature macrophage phenotype (Figure 2C) [Goh et al., 2023]. We next examined the expression of key markers associated with macrophage differentiation (Figure 2D). Consistent with previous reports, the fractalkine receptor *CX3CR1*, a marker of long-lived tissue-resident macrophages [Burgess et al., 2019], was most enriched in adult IEMs. Conversely, *CD34*, a common marker for HSCs and endothelial progenitors, was mainly expressed by the early FWs IEMs. *VEGFA* expression peaked in the middle FWs populations when the network of cochlear vasculature is increasing in density. The purinergic receptor *P2RY12*, implicated in regulating microglial surveillance and cAMP signaling, was enriched in both middle FWs and adult IEMs. As expected, *CSF1R*, which is essential for macrophage survival and homeostasis, was broadly expressed across all developmental stages. Interestingly, *FLT3* and *MYB*, key regulators of early hematopoiesis in bone marrow [Tsapogas et al., 2017, Wang et al., 2018], were exclusively expressed in IEMs aligned with the CMP/HSC trajectory. In contrast, *CDH5* (VE-cadherin), an endothelial marker suggestive of a hemogenic endothelium origin [Williamson et al., 2024], was expressed along the HLP trajectory. These IEM trajectory analyses were further interrogated using the Bian et al. [2020] dataset, and support a dual contribution to IEM seeding that is consistent with other organs (Supplementary Figure S5). Together, these results reveal distinct developmental pathways for macrophage ontogeny in the inner ear.

Independent of the developmental trajectories shown in Figure 2C, we observed a continuous spectrum of macrophage phenotypes spanning Mac 1, 2, 3, and 4 in order on the myeloid atlas (Figure 2E). Slingshot [Street et al., 2018] pseudotime analysis of the projected IEMs, using Mac 1 as the root, confirmed this spectrum, revealing a trajectory extending from Mac 1 to 4, while largely excluding Mac 6, 7, and most of Mac 5 (Figure 2F). When pseudotime was stratified by developmental age and subtype (Figure 2G), it showed no correlation with age but aligned strongly with subtype, indicating that this trajectory reflects a phenotypic transition independent of age-related differentiation. To further support this continuum, we examined smoothed gene expression patterns across macrophage subtypes (Figure 2H). The results revealed a gradual shift in gene expression from Mac 1 to 4, with Mac 2 and 3 sharing similar profiles and representing intermediate states.

### 2.3 Inner ear macrophages alternate their gene regulation profile during early development

Having investigated the influence of developmental age on macrophage subtypes, we next examined genes that were significantly up- or down-regulated in early FWs 7.5 and 9.2 compared to later developmental stages (middle FWs 16, 16.4 and adult; Figure 3A1). Early FWs IEMs showed enriched expression of *RTN4RL1* and *SEMA3C* (involved in the regulation of axonal outgrowth), *GABRE* (critical for GABA-A receptor production), and *NELL1* (implicated in osteoclast differentiation and bone formation). Subsequent gene set enrichment analysis (GSEA) of these age markers (Figure 3B1) revealed the potential involvement of IEMs in synaptic membrane development and limb morphogenesis, suggesting their broader roles in neural, skeletal, and tissue differentiation during early development. To identify transcriptional regulators of these age markers, we applied DecoupleR for transcription factor (TF) activity inference [Badia-i Mompel et al., 2022]. This analysis highlighted candidate regulators including *SMAD6* (a negative regulator of TGF-*β* signaling), *DACH1* and *FOXG1* (both important for nervous system development), *HDAC1* (involved in controlling cell proliferation and migration), as well as TFs associated with various macrophage functions including inflammation *MEIS1* and repair *ERG* and *GLI2* (Figure 3C1).

**Figure 3.**
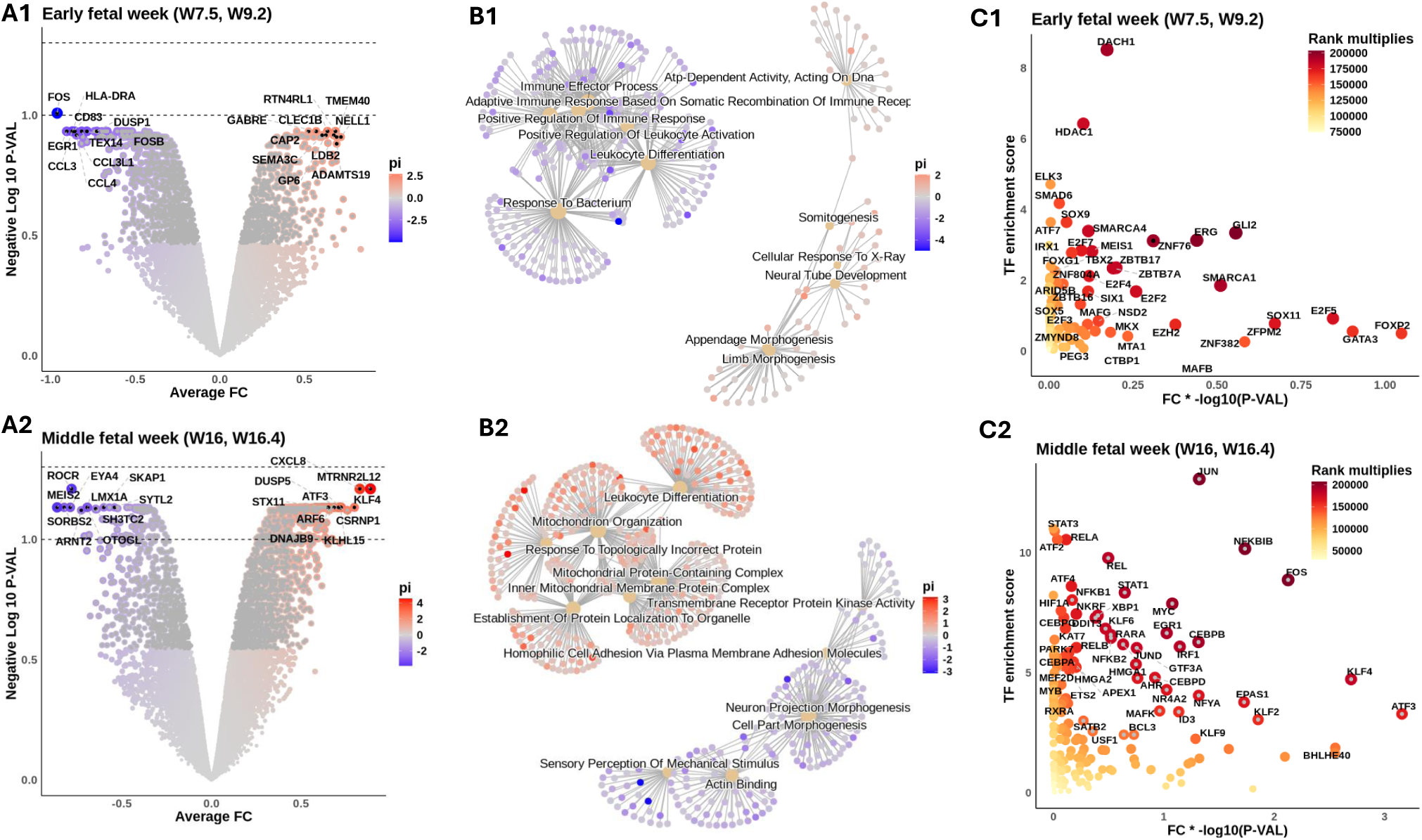
Intra- and intercellular signaling reveal macrophage functional heterogeneity during fetal development. Inner ear macrophages (IEMs) within each sample are aggreagted into a pseudobulk sample. Differential expression was conducted to compare fetal weeks (FWs) 7.5 and 9.2 samples to later developmental stages (middle FWs 16, 16.4 and the adult). Similarly, middle FWs samples are also compared to the other two age groups. The results for the early and the middle FWsare shown in subpanels numerated 1 and 2, respectively. **(A)** Volcano illustrating the DE results. Genes colored by its *π*-value = -log10(p-value) * fold change. Symbols of top 10 up- and downregulated genes are shown. Dashed lines show adjust p-value thresholds equal to 0.1 (lower) and 0.05 (upper) **(B)** Gene set enrichment analysis of age markers. The top 6 enriched terms for up- and downregulated genes are shown. Nodes connecting the terms are associated markers, colored by their *π*-values. **(C)** DecoupleR TF activity inference on DE results. x-axis: *π*-values of TFs. y-axis: DecoupleR activity scores of TFs. TFs colored by their multiplied ranking in *π*-values and activity scores. Symbols of top 50 ranked TFs are shown.

We then repeated these analyses for middle FWs 16 and 16.4, with comparisons made to all other donor ages (i.e., early FWs and adult; Figure 3A2). Middle FWs IEMs showed enriched expression of a broad range of immune-related genes, including *CXCL8* and *DUSP5* (involved in inflammation), *STX11* (regulating vesicle exocytosis), and *ARF6* (implicated in phagocytosis) (Figure 3A2). The enrichment of *KLF4*, *ATF3*, *CXCL8*, and *DUSP5* is intriguing, given their possible roles in pro-inflammatory functions. GSEA further highlighted the association of IEMs with inflammatory responses (Figure 3B2). The enrichment of mitochondrial activity suggests a metabolic switch at around FW 16, indicating a putative change in tissue microenvironment that occurs prior to the onset of hearing (Figure 3B2). Subsequent TF activity inference revealed a large number of immune-related TFs likely responsible for driving these transcriptional shifts from early to middle FWs. These TFs include regulators of innate immunity from the NF-*κ*B family (*NFKBIB*, *RELA*, *RELB*, *NFKB2*, and *NKRF*); key regulators of macrophage function including both pro-inflammatory (*ATF4*, *IRF1*, *TBP*, *XBP1*) and reparative (*STAT3*, *MYC*, *ERG1*, *FOS*) roles (Figure 3C2). In addition, *HIF1A* was highly enriched, providing further evidence for IEMs’ metabolic reprogramming around middle FWs.

### 2.4 Inner ear macrophages adopt distinct molecular identities in response to the dynamic tissue environment during fetal development

We applied NeighbourNet analysis to reconstruct gene regulatory networks (GRNs) of IEMs across developmental stages, characterising the dynamic of overarching regulatory patterns. This analysis prioritized the most significantly upregulated genes in each age group (identified by the DE analysis in Figure 3), predicted their regulatory interactions with TFs, and inferred upstream signaling cascades, starting from receptors that potentially transduce extracellular signals to regulate age marker expression through the predicted TF interactions. Complementing this, we employed NicheNet analysis [Browaeys et al., 2020] (Supplementary Results S1.1) to predict ligand sources, thereby tracing signaling origins and intercellular communication during HIE development.

At FW 7.5, the GRN prominently highlighted the upregulation of P-cadherin (*CDH3*), driven by *TGFBR2* –*SMAD3* and *FGFR1* –*CREBBP* signaling axes (Figure 4A1). This suggests a potential role for early fetal IEMs in maintaining sensory epithelial integrity and facilitating the proper formation of the tunnel of Corti via regulation of cell-cell adhesion [Beaulac and Munnamalai, 2024]. By FW 9.2, macrophages exhibited enriched SOX-family-mediated signalling involving *SOX5* and *SOX6*, which are TFs primarily known for their critical roles in chondrocyte and neuron differentiation (Figure 4A2) [Ji and Kim, 2016]. In our IEMs, these TFs are predicted to regulate chondro-osteogenic (*NELL1*) and neurogenic (*NELL1* and *SEMA3C*) targets. Hence, this observation suggests a macrophage-specific utilization of *SOX5/6* to support HIE development through coordinating otic-capsule ossification and guiding cochlear neural pathfinding.

**Figure 4.**
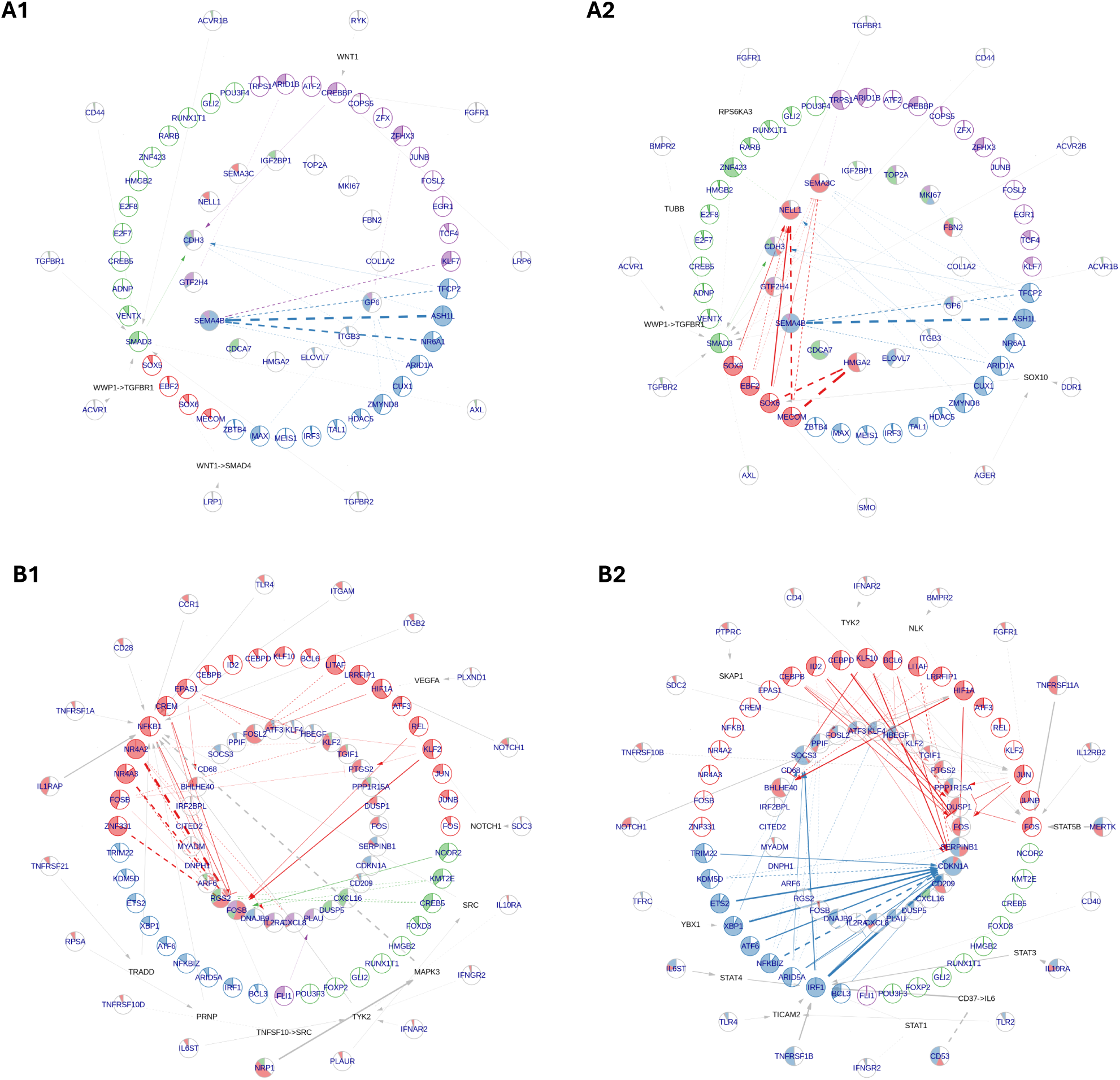
NeighbourNet inference of gene regulatory networks (GRNs) associated with the top 50 up-regulated genes by early and middle fetal week (FW) inner ear macrophages (IEMs). The age markers are derived from differential expression analysis analysisin Figure 3A. **(A)** GRNs for early FWs markers, constructed primarily by IEMs at (A1) FW 7.5 and (A2) FW 9.2. **(B)** GRNs for middle FWs markers, constructed primarily by (B1) Mac 2 and (B2) Mac 4 clusters. The macrophages used for GRN inference were automatically selected by the NeighbourNet algorithm as those exhibiting the most representative regulatory patterns. In each network, the innermost layer contains age markers, surrounded by their highly co-expressed transcription factors (TFs). Receptors are placed in the outermost layer, each connected to a TF predicted to mediate its regulatory influence. When the receptor–TF link is indirect, an additional layer displays the shortest inferred signalling path. Arrowheads indicate activation; bar-heads indicate repression; dashed lines denote links with significant co-expression but not supported by prior knowledge regulatory evidence.

The signaling landscape significantly shifted in macrophages during the middle FWs, adopting classical macrophage signaling pathways involved in immune and tissue modulation. Interestingly, major signaling variations within these macrophages were primarily driven by macrophage subtypes Mac 2 and Mac 4. Mac 2 displayed *NFKB1* -mediated pro-inflammatory signaling, relaying signals from IL1R (*IL1RAP*), TNFR1 (*TNFRSF1A*), CCR1, and TLR4 to upregulate FOS family genes, indicative of a transitional activation state (Figure 4B1) [Dorrington and Fraser, 2019]. In contrast, Mac 4 exhibited reduced NF-*κ*B signaling, instead activating *IRF1* -mediated pathways downstream of TNFR2 (*TNFRSF1B*), IL6R (*IL6ST*), and IL10R (*IL10RA*) to regulate key phagocytosis mediators *CDKN1A* and *SOCS3* (Figure 4B2) [Allouch et al., 2022, Gordon et al., 2016]. Furthermore, both macrophage subtypes demonstrated activation of a *NOTCH1* –*HIF1A* axis associated with macrophage responses to hypoxia and inflammation [Xu et al., 2015]. Notably, NicheNet analysis predicted that middle FWs IEMs were receiving pro-angiogenesis signals *VEGFA* (vascular endothelial growth factor A) (Supplementary Figure S6B). The activation of the *NOTCH* - *HIF1A* axis can, therefore, be explained by macrophage adaptation to transient hypoxic tissue niches created during vescular ingrowth in middle FWs, when local metabolic demand briefly outpaces the new circulation [Ramakrishnan et al., 2014]. Collectively, our findings reveal a broad spectrum of macrophage identities and functional roles shaped by the complex and dynamic microenvironment in the developing HIE.

## 3 Discussion

Recent single-cell studies have highlighted the heterogeneity of inner ear macrophages (IEMs) in mouse, revealing distinct subtypes that may contribute to the intricate developmental processes [Chiot et al., 2024]. Our data build upon these studies and histological investigations by [Steinacher et al., 2022] who report the presence of both resident and non-resident macrophages in the developing human inner ear (HIE) between gestational weeks 9 and 17.

Using a transcriptional approach, we have identified seven transcriptionally distinct IEM subtypes present over a similar window of HIE development (Figure 1D), with each subtype closely linked to a specific developmental age. For instance, we identifed a proliferative population of macrophages that infiltrate the early developing HIE (Mac 7), followed by the emergence of subtypes associated with neural development (Mac 3 to 6), including ion homeostasis (Mac 6), and subsequently subtypes with characteristics of mature antigen-presenting macrophages (Mac 1 and 2). Together, these results highlight, for the first time, the breadth of macrophage phenotypes present in the early developing HIE and their multidisciplinary contributions to normal development.

Moreover, lineage tracing studies in mice have provided insights into the developmental origins of IEMs, revealing their potential derivation from distinct embryonic sources [Kishimoto et al., 2019, Chiot et al., 2024]. Our data support these observations in human development, identifying inner ear seeding by both embryonic (yolk sac-derived) and more definitive (bone marrow-derived) macrophages during organogenesis. Specifically, we illustrate that FWs 7.5 to 9.2 IEMs cluster most closely with multipotent hemangioblasts (yolk sac progenitors), whereas FW 16 and adult IEMs show a hematopoietic stem cell origin based on the projection to the myeloid atlas (Figure 2C). These analyses are supported by additional projections of extensive human macrophage datasets [Bian et al., 2020] obtained across organs in human development (Supplementary Figure S4A,B,C and S5D,E). Moreover, presumptive yolk sac-derived IEMs (Mac 1 to 4) exhibit a distinct transitional trajectory that excludes Mac 5, 6 and 7 (Figure 2E). This result again supports multiple ontogenies of IEMs. When projected onto a human fetal macrophage atlas [Bian et al., 2020], IEMs display a unique tissue identity, aligning most closely with developing macrophages found in the skin and the brain (Supplementary Figure S4D,E,F). Our analyses support the conclusions from [Bian et al., 2020] that diverse macrophage subtypes are found at defined anatomical sites during human development. The precise contribution of each of these distinct macrophage lineages to their heterogeneity and functional diversity in HIE, is open for future exploration.

Our transcriptomic approach not only illuminates key developmental processes but also reveals new signaling interactions, with direct applications to macrophage-associated hearing pathologies [Pan et al., 2024, Sung et al., 2024]. Our data implicate numerous well-characterized trophic signaling pathways of macrophages, including TGF-*β*, FGF and semaphorin-neuropilin families in early developmental (FWs 7.5 and 9.2; Figure 4A, Supplementary Figure S6A), as well as *VEGFA*, *TNFSF*, *IGSF* and *CNTN2* during the middle developmental stage (FWs 16 and 16.4; Figure 4B, Supplementary Figure S6). In particular, the *TGFB2* and *VEGF* pathways are predicted to regulate *SEMA3A/C* expression in early fetal cochlear macrophages (Supplementary Figure S6A). Although the ligand *SEMA3A*, known to be important for normal cochlear morphology and function [Salehi et al., 2017], has traditionally been attributed to cochlear neurons and supporting cells [Cantu-Guerra et al., 2023], our analyses reveal that macrophages may be an additional cellular source.

We also identify a possible macrophage contribution to the newly discovered GABA signaling in the mammalian cochlea [Bachman et al., 2025], through their GABAA receptor subunit *GABRE* expression in early development (Supplementary Figure S6B). Further support for macrophage involvement in cochlear development is indicated by the enriched expression of *PDZRN3* in the Mac 6 population (Figure 1D), highlighting the potential role of this subtype in regulating Wnt signaling. Examining Wnt expression supports this hypothesis, illustrating Mac 6 as a source of Wnt5 ligands in HIE development (Supplementary Figure S3). Wnt signaling is critical for normal inner ear development [Munnamalai and Fekete, 2013], including *WNT5A* in correct hair cell function via planar cell polarity signaling [Dabdoub et al., 2003, Sewduth et al., 2014, Andre et al., 2012, Qian et al., 2007]. In addition, neuregulin signalling has been shown to be important for neural survival in the mammalian cochlea [Stankovic et al., 2004], and our analyses indicate Mac 6 as a source of both *NRG1* and *NRG3* in HIE development (Supplementary Figure S3).

Macrophages are also expected to contribute to the establishment of the intricate cochlear vasculature. The predicted high TF activity of *DACH1* in early FWs IEMs supports a critical role for macrophages in stria vascularis development (Figure 3C) [Ebrahim et al., 2016]. It is also possible that *DACH1* expression is under-represented in the middle FWs data, given that the stria vascularis is missing from these tissue dissections. *DACH1* is important for the development of endocochlear potential, with the knockdown of this TF causing hearing loss [Miwa et al., 2019]. Macrophages are therefore likely to play a pivotal role in normal cochlear development by orchestrating vascular formation, maintaining fluid homeostasis, and ultimately supporting the proper establishment of tonotopicity. These findings underscore the power of transcriptomic analyses in illuminating normal developmental processes, supporting the notion that IEMs play multidimensional roles far beyond their traditional immune functions. A deeper understanding of these diverse functions will not only enrich our fundamental knowledge of cochlear biology but also accelerate novel therapeutic strategies targeting immune-related, congenital, and age-related hearing loss [Pan et al., 2024, Sung et al., 2024].

### Limitations

A challenge in the present study is the limited availability of human donor tissue. Whilst we have captured early, middle, and late time points, we acknowledge that our conclusions are restricted to the tissues available. As further sequencing data become available, a more comprehensive spectrum of age-specific changes can complement this work, in addition to experimental animal studies to investigate fundamental biological questions.

Additionally, the paucity of human tissue has contributed to an imbalanced study design, in which macrophages from different age groups were collected from separate studies and dissected slightly differently (as noted). We acknowledge that frozen samples can yield variability in tissue quality. Consequently, comparisons among age groups may be confounded by batch effects and may not fully capture the variability present at each developmental stage. While we recognize that the multiple ontologies presented in Figure 2C would ideally be followed by proper pseudotime inference and de novo marker identification along pseudotime, the substantial technical differences between our self-sequenced-data and the dataset from van der Valk et al. [2023] preclude such integrative analyses.

## 4 Resource availability

The processed macrophage subset of the data, along with the code for reproducing all the bioinformatic analyses conducted, has been deposited in the Zenodo repository (https://zenodo.org/records/1

## 5 Acknowledgments

The authors would like to acknowledge the sample donors, the RCWIH BioBank, and the Princess Margaret Genomics Centre for their technical assistance. We thank the High Performance Computing Core for bioinformatics services, Compute Ontario (computeontario.ca) and the Digital Research Alliance of Canada (alliancecan.ca) for providing hardware storage and computing resources. We gratefully acknowledge constructive feedback on manuscript drafts from Drs Heiko Locher, Wouter van der Valk and Chao Wang.

## Funding

This study was supported by the Canadian Institutes of Health Research Scholarship for Doctoral Students and Master’s students; the Ontario Graduate Scholarship; the Raymond H. W. Ng Graduate Scholarship (BE); the Michael and Sonja Koerner Charitable Foundation (AD); the University of Melbourne - International collaborative grant scheme between Australia and Canada; the Australian Research Council (BN, LP190101139); Melbourne Research Scholarship (YD).

## 6 Author contributions

## 7 Declaration of interests

The authors declare that they have no competing interests.

## 8 Materials & Methods

The key resources used in this study, including the datasets analyzed and the softwares employed for the analysis are summarized in Table 1.

**Table 1.** This table summarizes the key resources used in this study, including the datasets analyzed and the softwares employed. **Resource name** indicates the name of the resource, **Source** refers to the publication or project from which the resource was generated, and **Identifier** provides the hyperlink for accessing the resource.

| Resource name | Source | Identifier |
| --- | --- | --- |
| **Data** |  |  |
| snRNA-seq human inner ear<br>Fetal weeks 16, 16.4 | Self-generated | https://zenodo.org/records/15328483/ |
| snRNA-seq human inner ear<br>Fetal weeks 7.5, 9.2 and adult | van der Valk et al. [2023] | GEO: GSE213796 |
| Myeloid cell atlas | Stemformatics & Rajab et al. [2021] | https://www.stemformatics.org/ |
| Macrophage module genes | Wang et al. [2023] | 10.1016/j.cell.2023.08.019 |
| **Software** |  |  |
| R v4.2.1 | The R project | https://www.r-project.org/ |
| Seurat v5.0.3 | Hao et al. [2024] | https://satijalab.org/ |
| scDblFinder v1.17.2 | Germain et al. [2022] | https://github.com/plger/scDblFinder |
| ggplot2 v3.5.0 | Wickham [2009] | https://ggplot2.tidyverse.org/ |
| clusterProfiler v4.11.1 | Yu et al. [2012] | https://bioconductor.org/packages/clusterProfiler/ |
| AUCell v1.18.1 | Aibar et al. [2017] | https://bioconductor.org/packages/AUCell/ |
| Sincast v1.0.0 | Deng et al. [2022] | https://github.com/meiosis97/Sincast |
| Slingshot v2.7.0 | Street et al. [2018] | https://bioconductor.org/packages/slingshot/ |
| limma v3.52.4 | Ritchie et al. [2015] | https://bioconductor.org/packages/limma/ |
| decoupleR v2.9.7 | Badia-i Mompel et al. [2022] | https://bioconductor.org/packages/decoupleR/ |
| nichenetr v2.2.0 | Browaeys et al. [2020] | https://github.com/saeyslab/nichenetr/ |

### 8.1 Ethics Approval

De-identified human fetal samples were obtained from the Research Centre for Women’s and Infants’ Health (RCWIH) BioBank with approval from the Research Ethics Boards of Mount Sinai Hospital (ID# 20-0003-E) and Sunnybrook Health Sciences Centre (Project Identification Number: 1514).

### 8.2 Human spiral ganglion collection and dissection

Samples were collected with written informed consent form between 2022 and 2023 from donors without known genetic or medical conditions. Three samples gestational week (GW) 18 were used in this study: a male and a female GW 18, and male GW 18.4. Sex was determined by PCR. Fetal gestational age was assigned via ultrasound (ACUSON Juniper Juniper^™^ Ultrasound System, Siemens Healthineers) by measuring the femur length, biparietal diameter, and foot length, then confirmed using a growth table [HERN, 1984]. These gestational ages have been converted into fetal weeks (FWs) to align with existing data used for comparison [van der Valk et al., 2023]. As such, GW 18 was included as FW 16, and GW 18.4 was included as FW 16.4. The time between collecting and receiving the tissue in our laboratory was less than 4 h. Samples were collected and dissected in ice-cold Hanks’ Balanced Salt Solution (HBSS) (Wisent, #311-512-CL) with 1% 1 M HEPES (Wisent, #330-050 EL). Upon receipt, inner ear tissues were isolated, and the spiral ganglia tissue were dissected. Two samples (female FW 16.0 and male FW 16.4) were treated for 10 minutes at 37^◦^C in 2 mg/mL thermolysin from Geobacillus stearothermophilus (Millipore-Sigma, #P1512) to decrease the amount of surrounding mesenchyme, dissected in the dissection solution containing Fetal Bovine Serum (ThermoFisher Scientific, #12484028) to immediately reduce the enzymatic activity and followed by washing steps with the solution mentioned above. Samples were placed in 1.5 mL Protein LoBind^®^ Tubes (Eppendorf, #022431081), then flash frozen and stored in liquid nitrogen.

### 8.3 Nuclei isolation & Sequencing

We modified the nuclei isolation protocol from 10x Genomics (Chromium Nuclei Isolation Kit, User Guide CG000505). Briefly, 500 *μ*L of lysis buffer were added to the tube containing the sample, incubated on ice for 1 min, and mechanically triturated with a P1000 pipette for up to 9 min and nuclei lysis was assessed throughout this process. Following cell lysis, nuclei were passed through a 40 *μ*m Flowmi^®^ Cell Strainer (Sigma-Aldrich). Then, centrifuged at 500 rcf for 3 min, and the pellet was resuspended in Debris Removal Buffer, and centrifuged at 700 rcf for 10 min. The pellet was then resuspended in Wash Buffer, centrifuged at 500 rcf for 10 min, and nuclei resuspended in Resuspension Buffer. To increase nuclei quantity, two out of the three samples followed an optimized protocol whereby after cell lysis, nuclei were washed in 500 *μ*L Wash Buffer and Resuspension Buffer (1:1). Nuclei quality was assessed throughout the protocol, and intactness was over 90% on average. The resuspended nuclei were loaded into the Chromium Chip (full capacity well) and processed following Chromium Next GEM Single Cell Multiome ATAC + Gene Expression workflow. cDNA libraries were sequenced using Illumina NovaSeq 600 and NovaSeq X. Raw BCL Illumina files were converted to FASTQ files using 10x Genomics Cell Ranger pipeline for demultiplexing and feature counting to generate gene expression matrices.

### 8.4 Bioinformatic analysis: data preprocessing

We analyzed two human inner ear (HIE) single-nucleus RNA sequencing (snRNA-seq) datasets in this study: one self-generated and another by van der Valk et al. [2023], which focuses on characterizing inner ear sensory development during fetal stages. Both datasets were preprocessed independently using the same pipeline described below.

We initiated preprocessing with the filtered unique molecular identifier (UMI) count matrices provided by the original studies, which were generated using the 10x Genomics Cell Ranger pipeline. To ensure data quality, we applied an initial quality control (QC) step to remove low-quality nuclei, including those expressing more than 8,000 genes, as well as those with mitochondrial transcript content exceeding 5%, indicative of potential cellular stress or ambient RNA contamination. Despite this initial QC, the total UMI distribution suggested the presence of doublets. Therefore, we applied doublet removal algorithm following cell clustering (will be described later on) to mitigate this issue.

We then applied the Seurat pipeline to normalize gene expression using log-normalization, with the median total UMI count as the scaling factor. Next, we identified the 2,000 most variable features (VFs) for each sample, scaled these VFs, and performed principal component analysis (PCA) [Hao et al., 2024]. Subsequently, we applied Seurat’s canonical correlation analysis (CCA)-based integration on the PCA space to harmonize the datasets, generating a lower-dimensional representation that preserves biologically coherent cell identities shared across samples [Butler et al., 2018].

Cell clustering was performed using Seurat’s shared nearest neighbor (SNN) - Leiden approach on the integrated CCA space to identify major cell populations. Cell clusters were then manually annotated based on marker genes identified through differential expression (DE) analysis using the MAST framework [Finak et al., 2015]. Finally, we applied scDblFinder to detect and remove doublets within each sample, leveraging the identified cell clusters to refine doublet classification. A final QC evaluation can be found in Supplementary Figure S7.

### 8.5 Identify human inner ear macrophage subtypes

From each preprocessed dataset, we subseted the macrophage population based on the expression of the marker genes *PTPRC* (CD45) and *ITGAM* (CD11b) (Supplementary Figure S1A2,B2). The selected macrophages were then combined into a single dataset. Following the same integration pipeline as applied to the full dataset, we reprocessed the macrophage subset using Seurat, performing VF selection, PCA, followed by CCA integration on the PCA space, treating samples as individual batches. Uniform manifold approximation and projection (UMAP) was applied to the PCA and the CCA space to visualize macrophage populations before and after integration, respectively.

We identified macrophage subtypes by performing SNN-Leiden clustering on the CCA space, setting the cluster resolution to 1, which resulted in seven clusters. This resolution was chosen to maximize the number of clusters while ensuring that each exhibited distinctly differentially expressed genes (DEGs) as revealed by MAST DE analysis [Finak et al., 2015]. A gene was considered DE in a cluster if it was expressed in at least 25% of the cells within the cluster, exhibited a log-fold change greater than 0.5, and had an adjusted p-value less than 0.05.

To functionally profile macrophage subtypes, over-representation analysis (ORA) of gene ontology (GO) terms associated with biological pathways was performed for DEGs in each cluster using ClusterProfiler [Yu et al., 2012]. GO terms with 10–500 genes were included, and those with an adjusted p-value below 0.05 were considered enriched. To remove redundancy and retain the most representative terms, we applied the simplify function in ClusterProfiler to enrichment results.

### 8.6 Profile inner ear macrophage identity by module gene expression

To characterize macrophage identity, we analyzed the expression of a predefined set of macrophage module genes, as identified by Wang et al. [2023]. These marker genes were derived from a comprehensive immune atlas of human fetal development, spanning multiple tissue types, and were used to classify primary macrophage subtypes. To explore how inner ear macrophages (IEMs) of different subtypes and ages express the module genes, we performed two key analyses: First, we conducted PCA on the expression of the module genes, generating a PCA space that is segregated by distinct modules. We projected macrophages onto this PCA space and visualized them alongside gene loadings using a PCA biplot. Second, we quantified the relative expression level of each module in individual cells using AUCell [Aibar et al., 2017], summarizing cellular identity as a vector of identity scores for each gene module.

### 8.7 Trace inner ear macrophage lineage by projecting onto an integrated myeloid atlas

To trace macrophage lineage, we queried our data against an integrated bulk gene expression atlas of myeloid cells. This atlas comprises samples from 44 independent studies, encompassing myeloid biology across diverse culture environments and various developmental stages. Here, ‘querying’ refers to projecting the query macrophage data onto the PCA space of the atlas. To achieve this, we applied the Sincast framework [Deng et al., 2022] to impute the single-cell query and align its distribution with the bulk atlas, enabling meaningful projection. The PCA of the atlas was performed on the 1,922 genes shared between the atlas and the top 15,000 VF of the query. The projection aligns the query cells with a stable gene expression space established by the atlas, highlighting biological differences while reducing technical noise. Therefore, leveraging the mapped query PC scores, we applied Slingshot [Street et al., 2018] to make robust estimation of pseudotime and cell lineage, with Mac 1 as the starting point for differentiation.

### 8.8 Reveal differences in inner ear macrophages signaling interactions during fetal development

We investigated macrophage signaling interaction unique to three age groups: early fetal weeks (FWs 7.5 and 9.2), middle fetal weeks (FWs 16 and 16.4), and adult (one sample). First, we performed DE analysis to pinpoint genes upregulated in each group. Next, we used curated prior knowledge databases to infer potential intra- and intercellular signaling interactions associated to these DEGs. For DE, we aggregated macrophages from each sample into pseudobulk profiles and rank-normalized their expression to reduce technical variability that may confound the analysis [Angel et al., 2020]. We performed DE to compare each age group with the two others using the limma pipeline, chosen for its suitability with limited sample sizes [Ritchie et al., 2015]. For each gene examined in a given age group, we calculated a *π*-value by multiplying the gene’s fold change by the negative log10 of its p-value. We used the *π*-value to determine the extent of DE [Xiao et al., 2014]. To functionally comprehend age differences, we performed gene set enrichment analysis (GSEA) on the resulting *π*-value–ordered list, focusing on GO biological pathway terms. Using ClusterProfiler, we tuned and refined GSEA as described for ORA on DEGs of macrophage subtypes.

To infer transcription factor (TF) activity representing intracellular gene regulation, we applied the DecoupleR algorithm [Badia-i Mompel et al., 2022] to each group’s *π*-values. This approach estimates TF activity by measuring the correlation between the TF’s known regulatory interactions (sourced from the CollectTRI database) and the observed *π*-values of target genes [Müller-Dott et al., 2023].

To extend beyond individual TF inference and elucidate overarching signaling dynamics across developmental stages, we employed NeighbourNet analysis [**?**] to reconstruct gene regulatory networks (GRNs) for age-specific marker genes (top 50 upregulated genes per age group based on *π*-values) and their potential upstream signaling pathways. For the markers of a given age group, NeighbourNet constructs GRNs at the level of individual cells and subsequently clusters and aggregates these networks to represent the principal gene regulatory patterns shared among cells. The top two aggregated GRNs for each of the early and middle FWs markers are displayed in Figure 4.

Finally, we utilized NicheNet to investigate intercellular signaling interactions, specifically ligand–receptor binding events implicated in gene upregulation within each age group [Browaeys et al., 2020]. For this analysis, we provided the identified age markers as target inputs. NicheNet then predicted upstream ligands, their corresponding receptors, and inferred their regulatory effects on the provided target genes.

## S1 Supplementary Results

### S1.1 NicheNet prioritization of ligands that target age-specific inner ear macrophage markers

We used NicheNet [Browaeys et al., 2020] to explore intercellular signaling between macrophages and other inner ear cell types during development. Specifically, we examined receptor-ligand interactions predicted to drive the upregulation of age markers derived from early FWs (Supplementary Figure S6A). Early FWs IEMs showed enriched signaling with chondrocytes (*CD74* -*COPA* and *NRP1* -*SEMA3C/3D*), cochlear epithelium (*TGFBR1/2/3* -*TGFB2* and *FZD2/6* -*SFRP1*), melanocytes (*TGFBR1/2/3* -*TGFB2* and *GJB2* -*GJB6*), neurons (*FGFR1/2* -*FGF10* and *NRP1/2* - *SEMA3E*), and IEMs themselves (*PDGFRA/B*, *PLXNA1* -*NRP1*, and *TGFBR1/2/3* -*TGFB1*). These interactions highlight an early activation of growth factor pathways *TGFB1/2* and *FGF1/10*, which are predicted to upregulate genes such as *HMOX1*, *TOP2A*, *SEMA3A/3C*, *HMGA2*, and *ADAMTS19*. Collectively, these results support a previously unidentified trophic role for early IEMs during development (Figure 1D). Notably, some of the TGF-*β* and FGF targets are also implicated in neural outgrowth and pathfinding.

A similar NicheNet analysis was applied to age markers derived from middle FWs (Supplementary Figure S6B). In comparison, the middle FWs IEMs were predicted to adopt a classical macrophage phenotype and predominantly communicate with mesenchymal, endothelial, and other IEMs. It should be reiterated that the tissues collected at this donor age were isolated exclusively from the cochlear modiolus. Therefore, these data should be interpreted within the context of this specific inner ear location. As such, the cell populations shown in Supplementary Figure 1A1 were enriched; however, cochlear epithelial cells (including hair and supporting cells), as well as cells of the stria vascularis and lateral cochlear wall, were absent. Our analyses reveal that modiolar macrophages were predicted to communicate with ACAN + mesenchymal stem cells (*IGSF11* -*IGSF*), endothelial cells (*TNFSFRSF10A/B/C/D/11B* -*TNFSF10*, *ERBB2* -*HLA-A*, *LILRB1/2* -*HLA-A*, and *KLRB1/KLRF1* -*CLEC2B*), and modiolar macrophages themselves (*CD4/9/37/53/63/81/82* -*HLA-DRA*). Collectively, these predicted signaling interactions suggest age-dependent regulation mediated by *VEGFA* and *TNFSF10* family ligands, targeting genes essential for core macrophage functions, including phagocytosis (*CDKN1A*), wound healing (*HBEGF* and *PLAU*), and efferocytosis (*ANXA1*). Interestingly, the predicted target genes also spanned both known inflammatory (*BHLHE40*, *PPP1R15A*, *PTGS2*, *IL2RA*, and *SOCS3*) and reparative (*CXCL8*, *DUSP1*, *KLF2*, *KLF4*, *MAFF*, *PLAU*, *ATF3*, *SOCS3*, *PPIF*, *CITED2*, *FOSL2*, *NFIL3*, and *RGS2*) immune response programs, reflecting a possible surveillant or transitional activation state in modiolar macrophages at this stage.

## S2 Supplementary Figures

**Figure S1.**
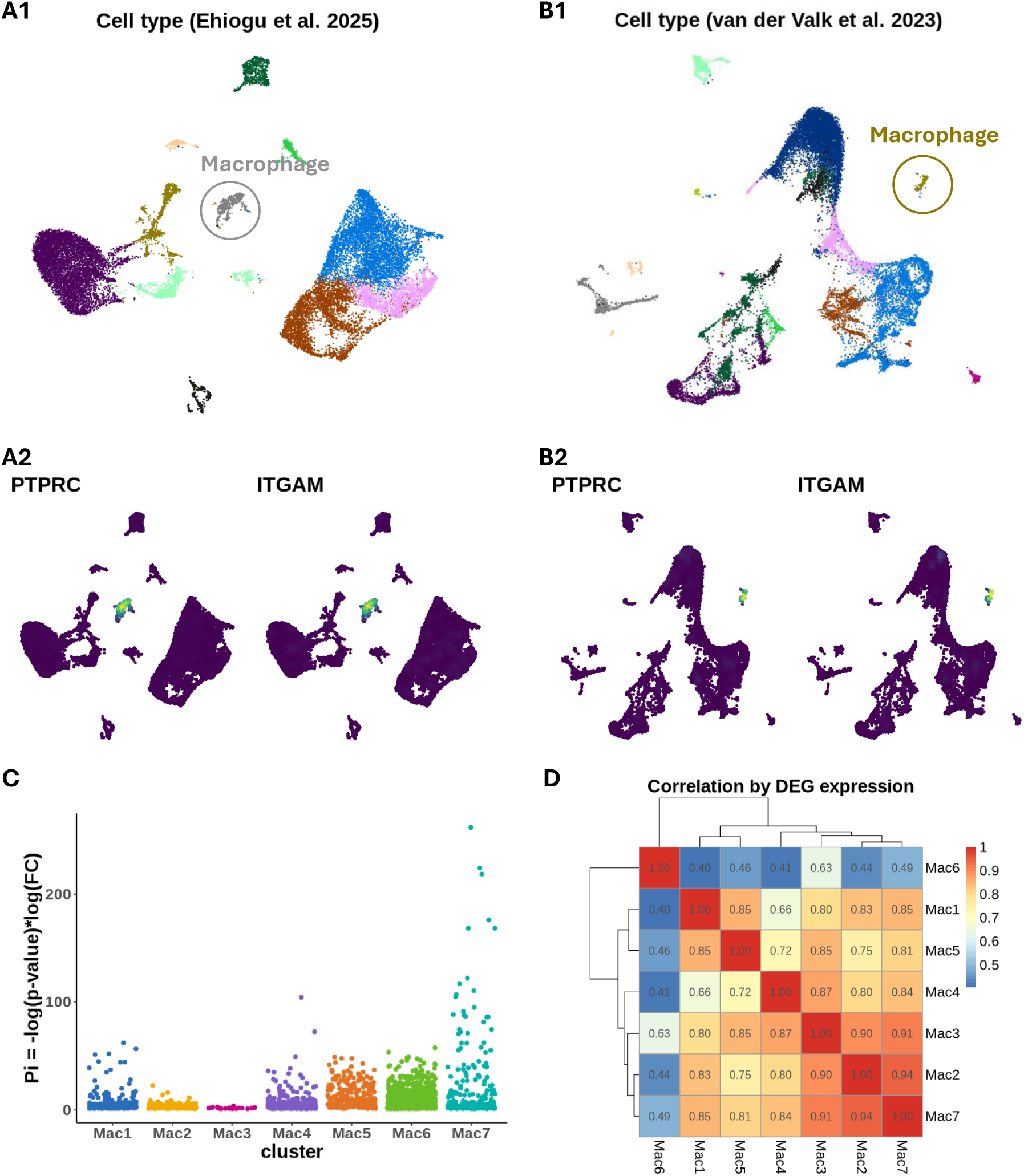
Selection of macrophage subsets from human inner ear snRNA-seq data. **(A)** UMAP plots showing the distribution of cell types from Ehiogu et al. (2025). A2: Nebulosa highlights the density of expression of two macrophage markers used for macrophage selection: PTPRC (CD45) and ITGAM (CD11b). **(B)** Similar to (A), but showing the UMAP plots of van der Valk et al. [2023]. **(C)** *π*-value (y-axis: negative log p-value × log fold change) indicates the level of differential expression of DEGs, identified for each macrophage subtype (x-axis). **(D)** Correlation matrix displaying the relationship between different macrophage subtypes’ pseudobulk expression profiles in DEGs.

**Figure S2.**
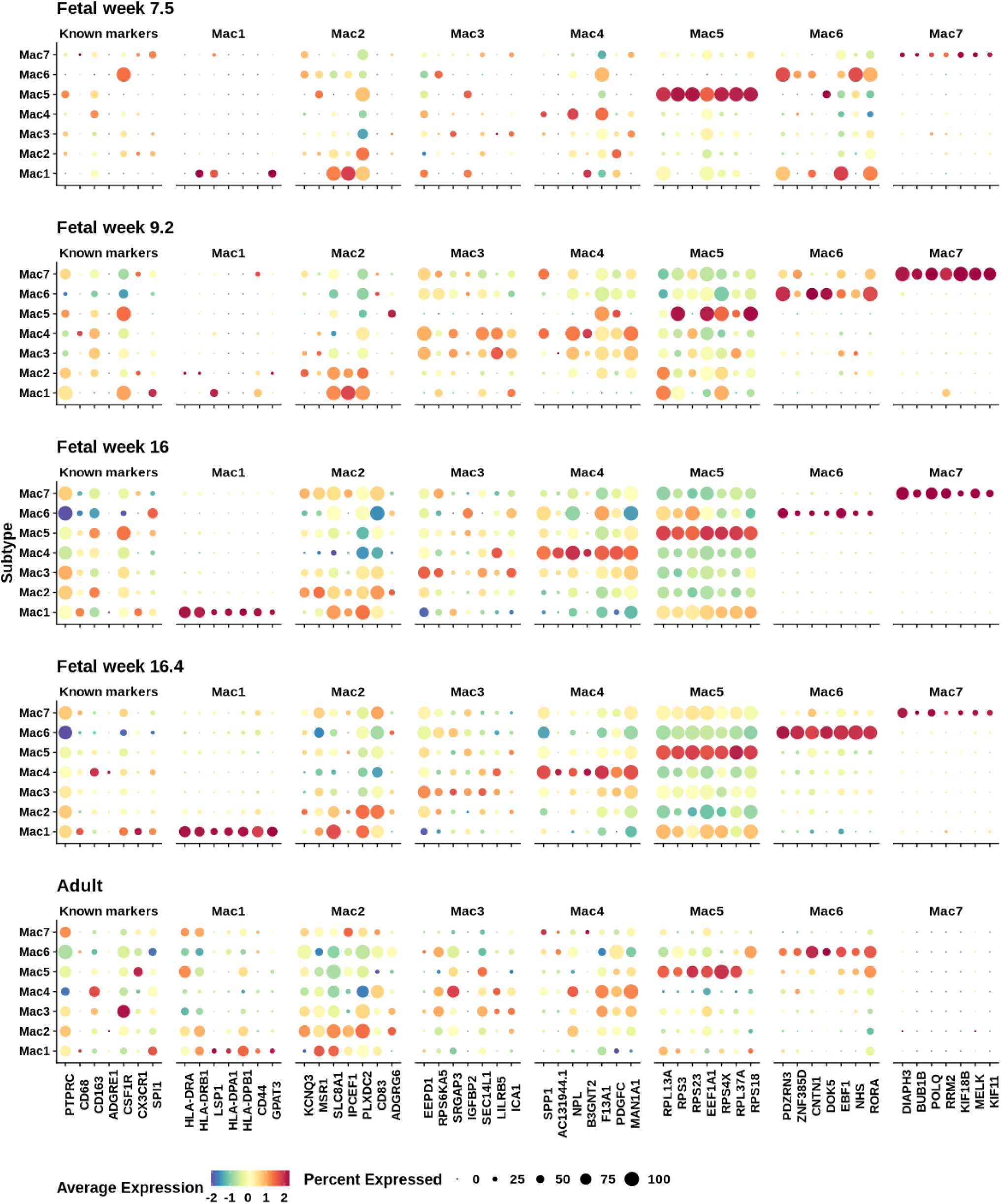
Macrophage marker genes. Expression of common macrophage markers, and top ranked discriminating genes (first 7 genes) in each subtype. In addition to showing the expression in each age group independently to demonstrate the consistency of markers across age groups, the figure notation follows the same style as Figure 1.

**Figure S3.**
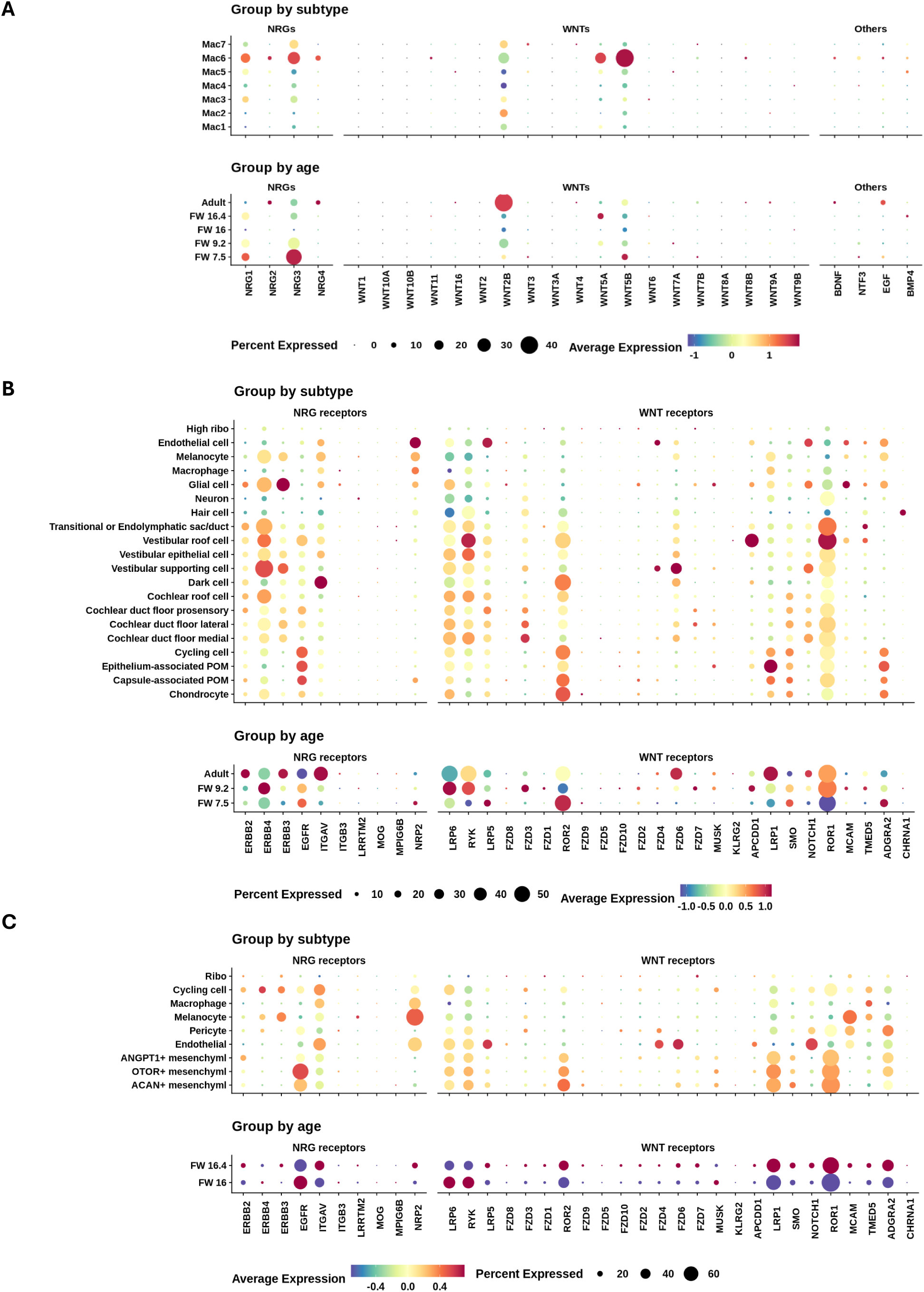
Growth factor signaling. Interpreted as in Figure 1D, this figure profiles growth-factor and receptor expression related to NRG and WNT signaling across inner-ear cell types. **(A)** Growth-factor (ligand) expression in macrophages. **(B)** and **(C)** WNT and NRG receptor expression in early fetal week (Week 7.5 and 9.2) and middle

**Figure S4.**
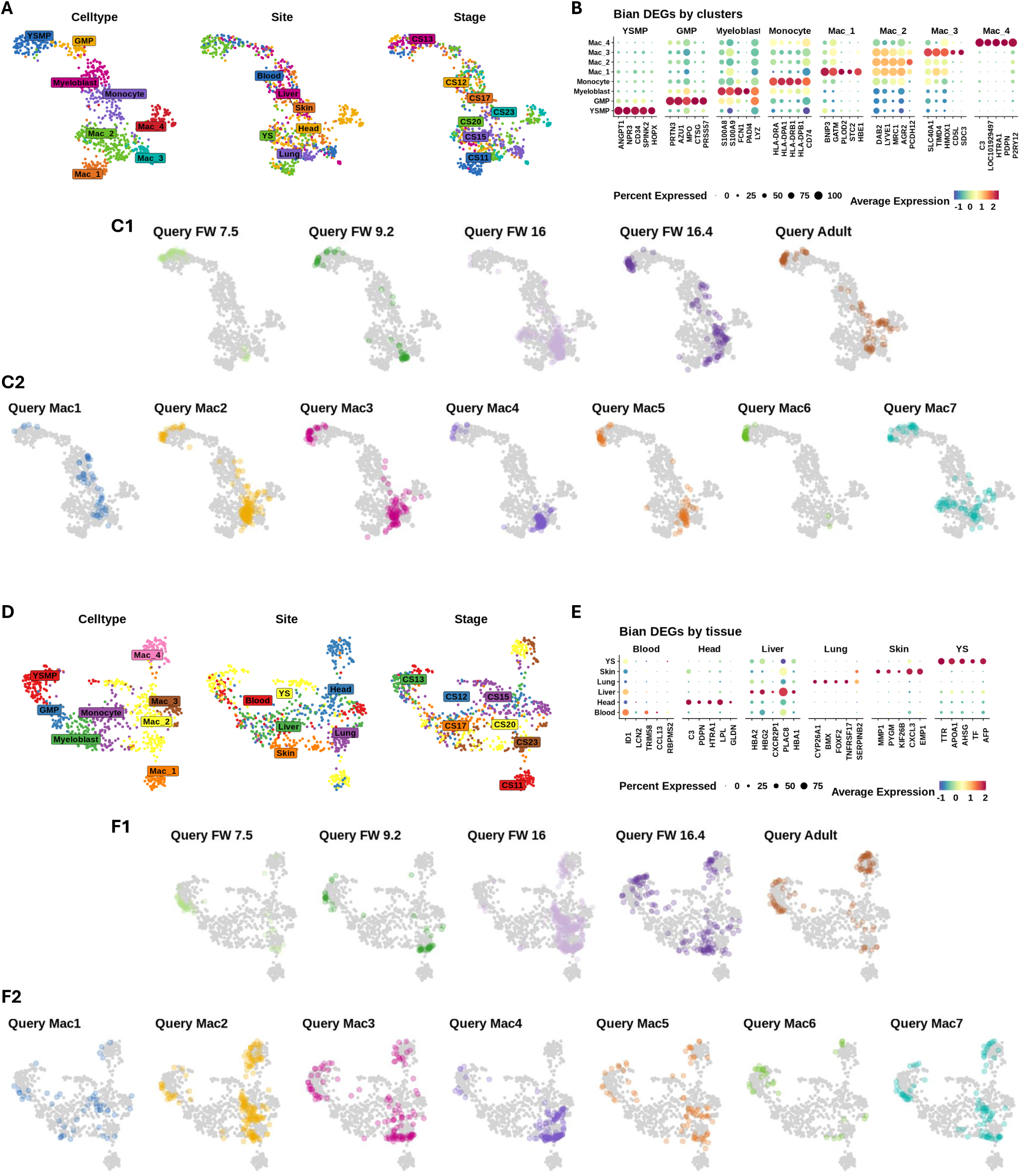
Benchmarking inner ear macrophage identity using Data from Bian et al. [2020]. **(A)** UMAP embedding of the Bian et al. [2020] dataset, based on the 2,000 most variable features. Cells are colored by their assigned cell type, tissue of origin, and developmental stage. **(B)** Expression of the first five celltype markers. Celltypes are shown on the y-axis, and gene symbols on the x-axis. The figure notation follows the same style as Figure 1. **(C)** Projection of query IEMs onto the UMAP in (A). Panels (C1) and (C2) show query cells colored by age and by macrophage subtype, respectively. **(D)** UMAP embedding of the Bian et al. [2020] dataset based on upregulated tissue-specific markers. **(E)** Similar to (B), but showing expression of the first five tissue-specific markers. **(F)** Similar to (C), with the query IEMs projected onto the UMAP in (D).

**Figure S5.**
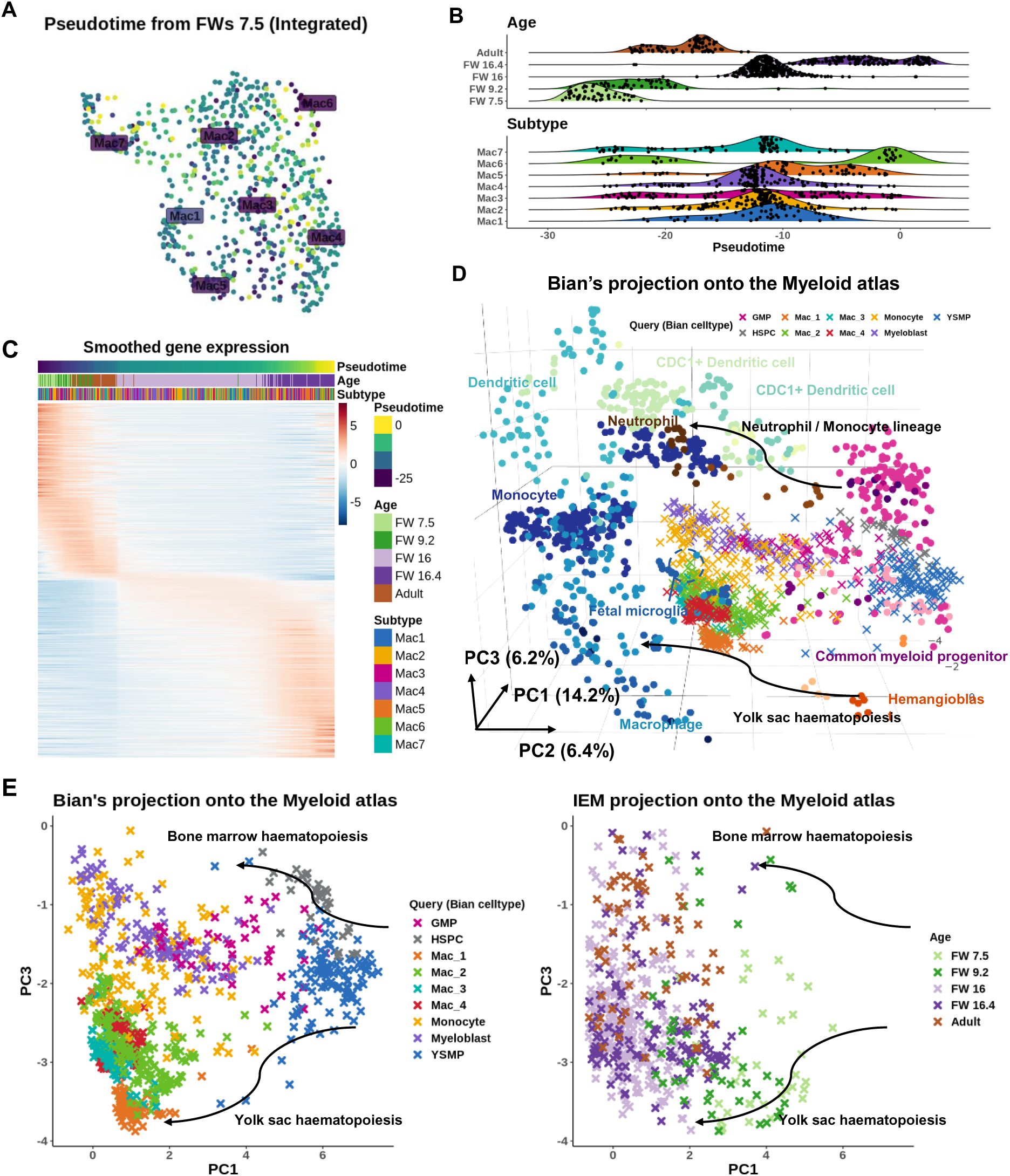
Extended trajectory analysis. (A-C) are similar to Figure 2(F-H), but pseudotime is recalculated in Slingshot using fetal week (FW) 7.5 macrophages as the root. **(D)** Mirrors figure 2(C), but showing the projecting the fetal macrophage data from Bian et al. [2020] onto the reference myeloid atlas. **(E)** Side-by-side comparison of Bian’s projection (L) and the IEM projection (R) reveals that early inner ear macrophages (FWs 7.5 and 9.2) overlap with the yolk-sac macrophage lineage present in the Bian’s study.

**Figure S6.**
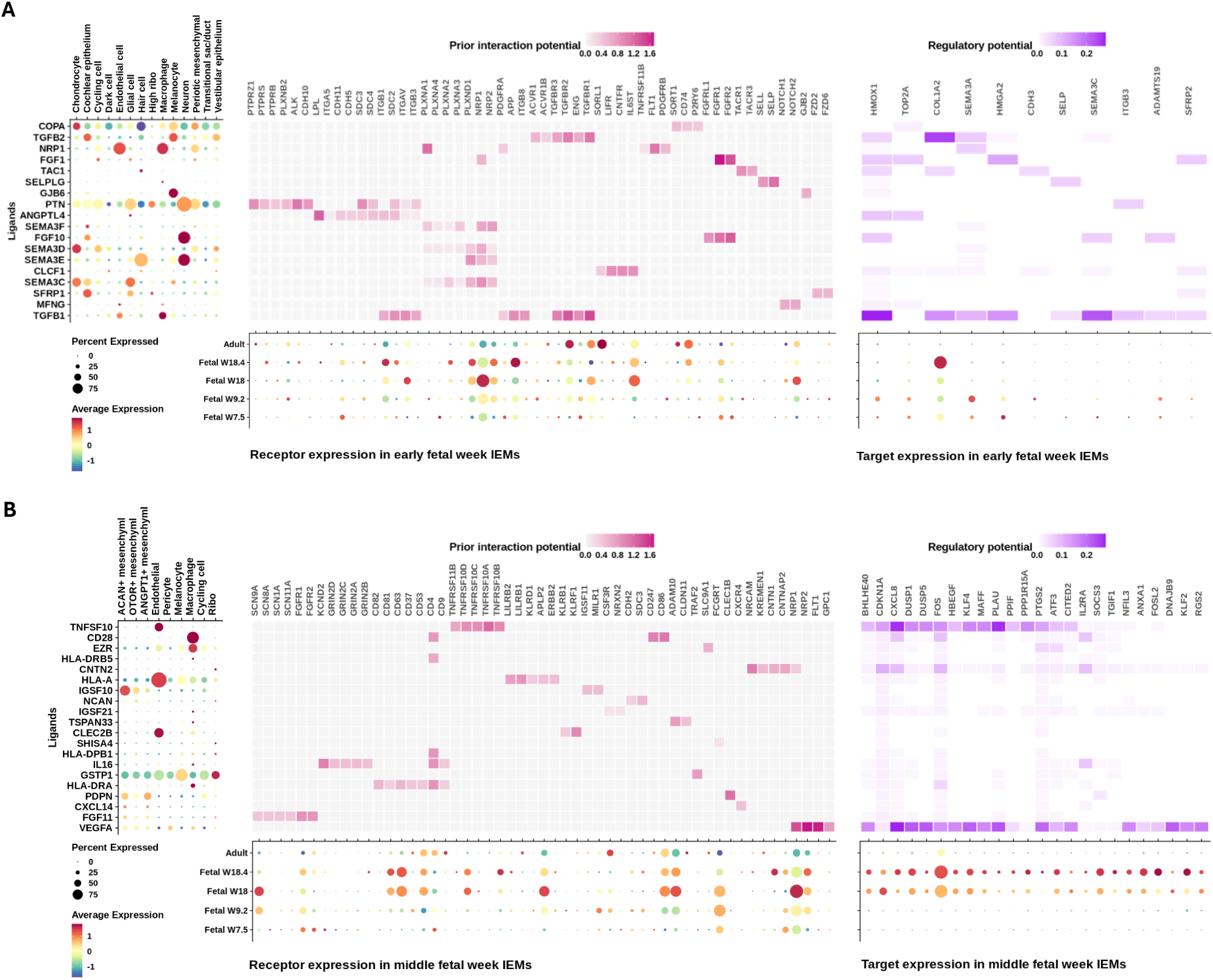
Prioritizing ligands with high regulatory potential on the top 50 age markers using NicheNet. Age markers are the upregulated genes of A age group derived from the differential expression analysis in Figure 3A. **(A)** NicheNet ligand prioritization using the age markers of early fetal weeks (FWs 7.5 and 9.2) inner ear macrophages (IEMs) as the targets of interests. **(B)** Similar to (A), but showing NicheNet analysis on the age markers of middle FWs (FWs 16 and 16.4) IEMs. Pink heatmap: NicheNet ligand (row) - receptor (column) interaction weights. Purple heatmap: NicheNet ligands (row) on targets (column) regulatory potential. Aligned to the rows of the heatmaps is the dotplot showing the expression of the prioritized ligands across inner ear cell types. Aligned to the columns of the heatmaps are the dotplots showing the expression of predicted receptors and targets within IEMs.

**Figure S7.**
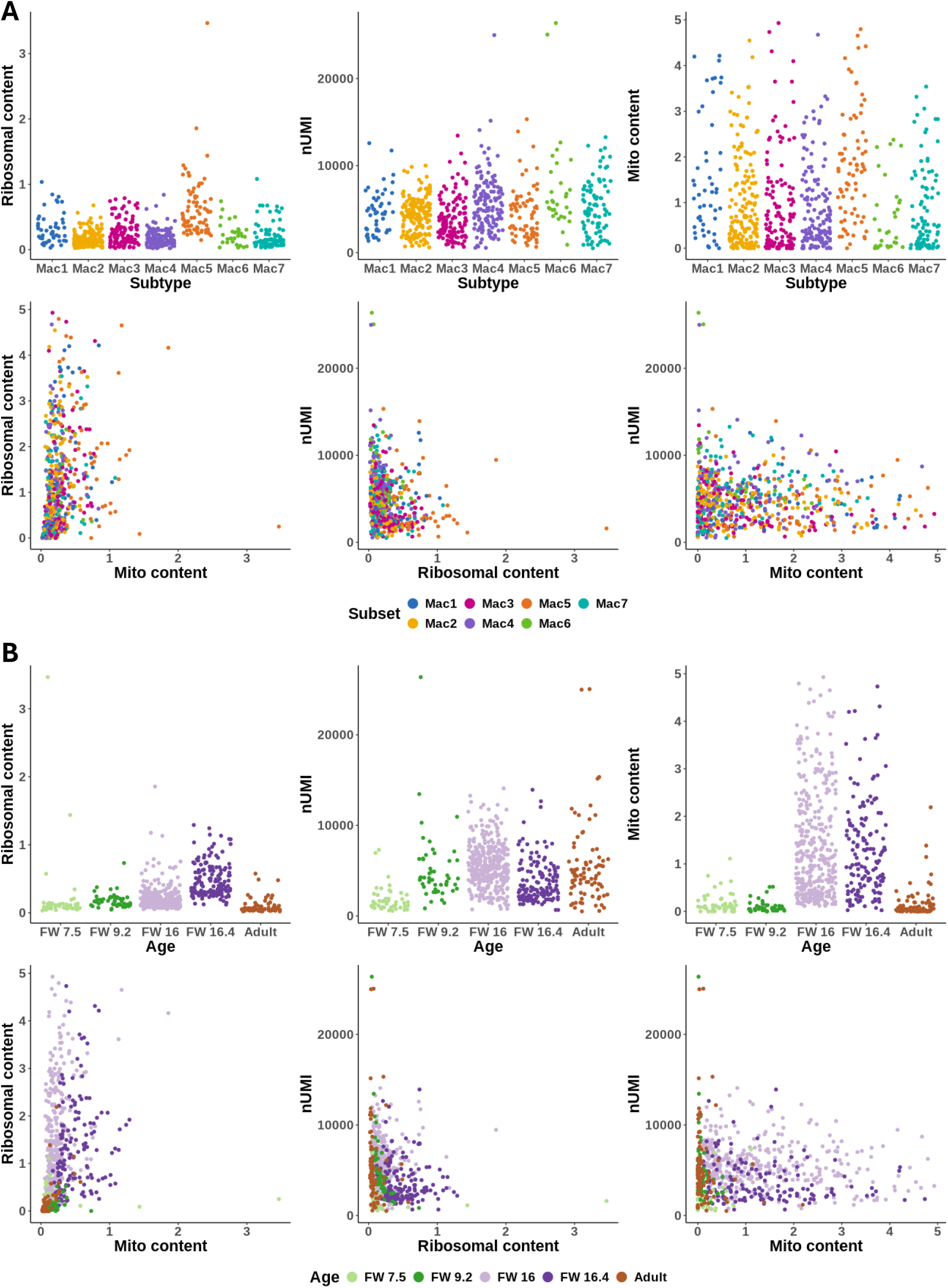
Quality checking of inner ear macrophages. Quality checking of IEMs by assessing ribosomal content, mitochondrial content, and total UMI counts. Macrophages are grouped by (A) subtype and (B) age. No strong association is observed between any QC metric and macrophage grouping, suggesting overall high data quality after pre-processing.

## Notes

https://zenodo.org/records/15328483/

### Competing Interest Statement

The authors have declared no competing interest.

### Summary of Updates

This version of the manuscript has been revised to include new results.

